# Aging amplifies the influence of spatial contextual information on visual scene processing

**DOI:** 10.64898/2026.02.20.706940

**Authors:** Clément Naveilhan, Raphaël Zory, Stephen Ramanoël

**Affiliations:** Université Côte d’Azur, LAMHESS, Nice, France; Institut Universitaire de France (IUF), Paris, France; Sorbonne Université, INSERM, CNRS, Institut de la Vision, 17 rue Moreau, F-75012 Paris, France

**Author notes:** CA, Corresponding Author: Clément Naveilhan & Stephen Ramanoël.

**Keywords:** Aging, Visual perception, Spatial memory, Prior information interaction

## Abstract

Older adults rely increasingly on prior knowledge to make sense of their deteriorating representation of the visual world, but how this shapes scene perception and spatial reorientation remains unclear. To address this issue, 28 young and 25 older adults viewed artificially generated rooms either before or after learning the position of a goal hidden in an adjacent room. We manipulated both the number and the eccentricity of navigational affordances (*i.e.,* open doors) to investigate the interaction between bottom-up scene features and top-down spatial knowledge. Consistent with previous findings, younger adults showed decreased performance as the number of open doors increased, but only after learning the goal’s position, indicating a top-down interaction with the automatic processing of affordances. Door eccentricity did not affect this interaction, suggesting our findings were not due to a distractor effect. In older adults, this interaction between prior spatial information and navigational affordances was markedly amplified: reaction times increased at twice the rate observed in younger adults. These findings show that prior spatial knowledge interacts with the automatic extraction of navigational affordances, and that this influence is markedly amplified with age. While prior knowledge helps stabilize perception when sensory processing becomes less reliable, it can also increase the processing time for complex scenes, particularly when multiple action possibilities are present. By revealing how aging shifts the balance between top-down and bottom-up mechanisms, these results refine models of age-related spatial navigation decline and highlight a trade-off whereby increased reliance on prior knowledge supports perception but can also slow interaction with complex environments.

## Introduction

Spatial navigation is a fundamental cognitive ability that enables individuals to orient in space, plan routes, and reach goals efficiently. It is also among the functions most vulnerable to aging and has been proposed as an early clinical marker of cognitive impairment (Coughlan et al., 2018; Lester et al., 2017; Lithfous et al., 2013; Moffat, 2009). While age-related declines in spatial navigation have traditionally been attributed to impairments in memory or executive functions, growing evidence indicates that visuospatial processing plays a central role in navigational performance (Ekstrom, 2015). This processing does not operate in isolation, but is dynamically coupled with top-down knowledge, shaping how navigational information is selected, interpreted, and used (Qi & Mou, 2024). With aging, however, this tightly coupled system is further constrained by age-related changes in basic visual functions, such as acuity or contrast sensitivity, which alter the processing of visual scene features and ultimately impair spatial reorientation (Durteste et al., 2023; Naveilhan et al., 2025; Owsley, 2016; Ramanoël et al., 2015, 2020). Yet, how impairments in bottom-up information processing during aging interact with the well-documented decline in higher-order cognitive functions remains poorly understood. Addressing this gap is essential to move beyond accounts that emphasize cognitive factors alone in age-related navigational deficits and to clarify how changes in perceptual and memory processes jointly contribute to spatial navigation impairments (Pepe et al., 2025).

As humans, our capacity to navigate relies mainly on visual information, enabling the rapid extraction and effective use of rich visuospatial cues. Scene perception provides the basis for spatial orientation by encoding the structural layout of the environment, thereby supporting cognitive map formation and guiding goal-directed behavior (Epstein & Baker, 2019; Julian et al., 2018). The interplay between visual perception and navigation is supported by three scene-selective brain regions: the parahippocampal place area (Epstein & Kanwisher, 1998), involved in scene categorization and landmark processing; the medial place area (Silson et al., 2016), witch encodes visual scene layout cues to compute world-referenced orientation; and the occipital place area (Dilks et al., 2013), which encodes egocentric distance and local scene elements to guide first person vision-based navigation. Neuroimaging studies have recently extended this framework by identifying three adjacent place memory areas that form a continuous posterior–anterior gradient from visually driven scene representations to memory-based spatial representations (Steel et al., 2021, 2023, 2024). In the context of aging, this network undergoes several modifications (Ramanoël et al., 2019, 2020), with a functional dedifferentiation and loss of neural selectivity resulting in a deterioration of fine-grained spatial processing, observable in the age-related decline of several visual functions (Koen & Rugg, 2019; Ramanoël et al., 2015).

Thus, the age-related reduction in the fidelity of bottom-up sensory signals has direct functional consequences, leading to a systematic reweighting of information sources under uncertainty for older adults. That is, as the visual inputs deteriorate with age, perception relies more strongly on priors and contextual expectations (*i.e.,* top-down modulation) to resolve ambiguity (Chan et al., 2021; Gilbert & Moran, 2016; Hsu et al., 2021; Tarasi et al., 2026; Trapp et al., 2022). Such reweighting is especially apparent under noisy conditions, where older adults can show greater benefits from top-down guidance, aligning with broader accounts of age-related changes in the neural systems that support visual perception and attention (Madden et al., 2004; Whiting et al., 2014). However, this is not uniformly beneficial across tasks, and older adults can be less efficient at using explicit top-down modulation to constrain visual sampling during search (Barrett et al., 2024). Similarly, related work indicates age-related differences in how contextual versus sensory information is weighted during prediction under uncertainty (Ravizzotti et al., 2025). These results converge to indicate a compensatory mechanism wherein older adults increasingly depend on top-down modulation based on semantic cues to interpret noisier bottom-up sensory input. Recently, Merhav et al. (2019) extended these findings to navigation, proposing that an increased persistence of previously experienced spatial contexts can hinder spatial updating in older adults, thereby offering a mechanistic account of the commonly observed age-related decline in spatial memory. In this view, older adults may rely more heavily on top-down modulation, such as schema-driven expectations about scene structure (Peelen et al., 2024), to compensate for degraded sensory input. However, this compensatory reliance can also incur performance costs when reinstated information persists despite no longer being accurate or task-relevant (Wynn et al., 2020), an apparent contradiction that highlights the critical need to delineate this interplay.

Disentangling the respective contributions of bottom-up and top-down modulation in visual cognition paradigms remains a methodological challenge. In particular, it requires the use of comparable visual stimuli and tasks that engage processing of the same task-relevant features, while allowing for systematic manipulations of the availability of scene information and prior knowledge. In this context, navigational affordances (*i.e.,* action possibilities offered by the visual layout of a scene) provide a useful conceptual and experimental framework (Bartnik et al., 2025; Djebbara et al., 2019; Gregorians & Spiers, 2022; Naveilhan et al., 2024). While early studies emphasized automatic extraction of affordances through bottom-up visual processing (Bonner & Epstein, 2017; Harel et al., 2022), growing evidence highlights a significant role for top-down modulation of affordance processing apparent at both behavioral and neural levels (Aminoff & Tarr, 2021; Choi et al., 2020; Naveilhan et al., 2024; Rossel et al., 2022). For example, in previous work from our group with young adults, we showed that learning the spatial location of a goal to be retrieved interacts with the bottom-up extraction of scene features, such as the number of available open doors within a room (Naveilhan et al., 2024). These behavioral effects were supported by electroencephalographic evidence indicating that prior spatial knowledge immediatly modulates early stages of visual scene processing (Naveilhan et al., 2025). Taken together, these studies underscore the value of the navigational affordances framework to investigate the interaction between prior spatial information and bottom-up processing. As previous work has highlighted, understanding how these interactions are impacted by aging is particularly crucial, as older adults increasingly rely on top-down modulation to compensate for degraded bottom-up information and such insights may ultimately guide interventions to support navigation in older populations (Sheynikhovich et al., 2025).

To this end, we adapted a navigational task from previous work (Naveilhan et al., 2024) and compared behavioral performance between young and older adults. Participants were presented with artificially generated rooms on a screen, either before or after learning the position of a goal they had to reach in an adjacent room. Across both experimental phases, we manipulated the number of navigational affordances (open doors) to examine the interaction between bottom-up processing of scene features and prior spatial information. In addition, we varied the eccentricity of lateral affordances to control for potential distractor effects, motivated by prior evidence showing that distractor interference scales with eccentricity (Chen, 2008; Chen & Treisman, 2008; Honda, 2005). First, we aimed to replicate previous findings in younger adults, namely a decrease in performance as the number of affordances increased, but only after participants had acquired the spatial information of goal position. Second, we expected that eccentricity of the affordances would not interfere with the extraction of the number of navigational affordances, arguing against the interpretation that additional doors act only as visual distractors. Finally, while a general age-related increase in reaction time is expected, current literature suggests that older adults increasingly rely on top-down modulatory processes. Based on this framework, we can posit two mechanistic accounts that yield dissociable behavioral predictions. First, as previous studies indicate that older adults exhibit degraded processing of low-level scene features and a greater dependence on contextual cues, they may not show modulation by the number of affordances, instead relying only on prior contextual information. Alternatively, the increased age-related reliance on prior information may enhance the interference with affordance extraction, leading to a greater reaction time cost with increasing number of affordances after the goal location is learned, amplifying the effect beyond what is observed for younger adults. Disentangling these two possibilities is critical for determining whether age-related navigation deficits are mitigated by enriched contextual information or, conversely, whether such information interferes with the processing of scene-based features and further degrades performance.

## Method

### Participants

The present behavioral experiment involved 28 young adults (23.29 ± 2.94 years old; range 19 – 29; 13 females) and 25 healthy older adults (73.63 ± 6.57 years old; range 64 – 85; 14 females). The required sample size was determined using G*Power (version 3.1; Faul et al., 2007), based on effect sizes observed in a previous study (Naveilhan et al., 2024). To detect an effect size of *d* = 0.76 with an alpha level of 0.05 and a statistical power of 0.95 in a between-subjects design, a total of 46 participants (23 per age group) was needed. All participants reported normal or corrected-to-normal vision and had no history of neurological or psychiatric disorders. Neurocognitive status in older adults was screened using the Montreal Cognitive Assessment (MoCA; Nasreddine et al., 2005), with all older participants scoring above the threshold of 26 (mean 27.56 ± 1.41). The study received approval from the local ethics committee (CERNI-UCA, number 2021–050), and all participants provided written informed consent prior to participation.

### Stimuli and procedure

The visual stimuli were created using the Unity Engine software (version 2019.2.0.0f1) and consisted of minimalistic rectangular virtual rooms. Each room included either a door or a gray rectangle on each of the three visible walls to maintain consistent visual complexity across conditions (Bonner & Epstein, 2017). We manipulated the eccentricity of the lateral walls, thereby modifying the spatial position of lateral navigational affordances and gray rectangles (**Figure 1.A**). In the *narrow condition*, the center of each lateral element was positioned 11.5 cm from the center of the screen, corresponding to approximately 11° of visual angle on each side. In the *control condition* affordances or gray rectangle were positioned at 17 cm (16.22° of visual angle), and in the *wide condition*, at 22.5 cm (21.28° of visual angle). This manipulation allowed us to systematically vary the perceptual eccentricity of the lateral affordances while preserving the overall scene structure and maintaining a constant central door position. We did this to control potential attentional confounds, since distractor interference is known to vary with eccentricity (Chen, 2008; Chen & Treisman, 2008; Honda, 2005). Accordingly, if the observed interaction between the number of affordances and prior information persists across *narrow*, *control*, and *wide* conditions, this would indicate that the effect cannot be explained by a purely distractor-based account. On the other hand, if affordances are considered as distractors, we would report a modulation of performances with eccentricity.

**Figure 1.**
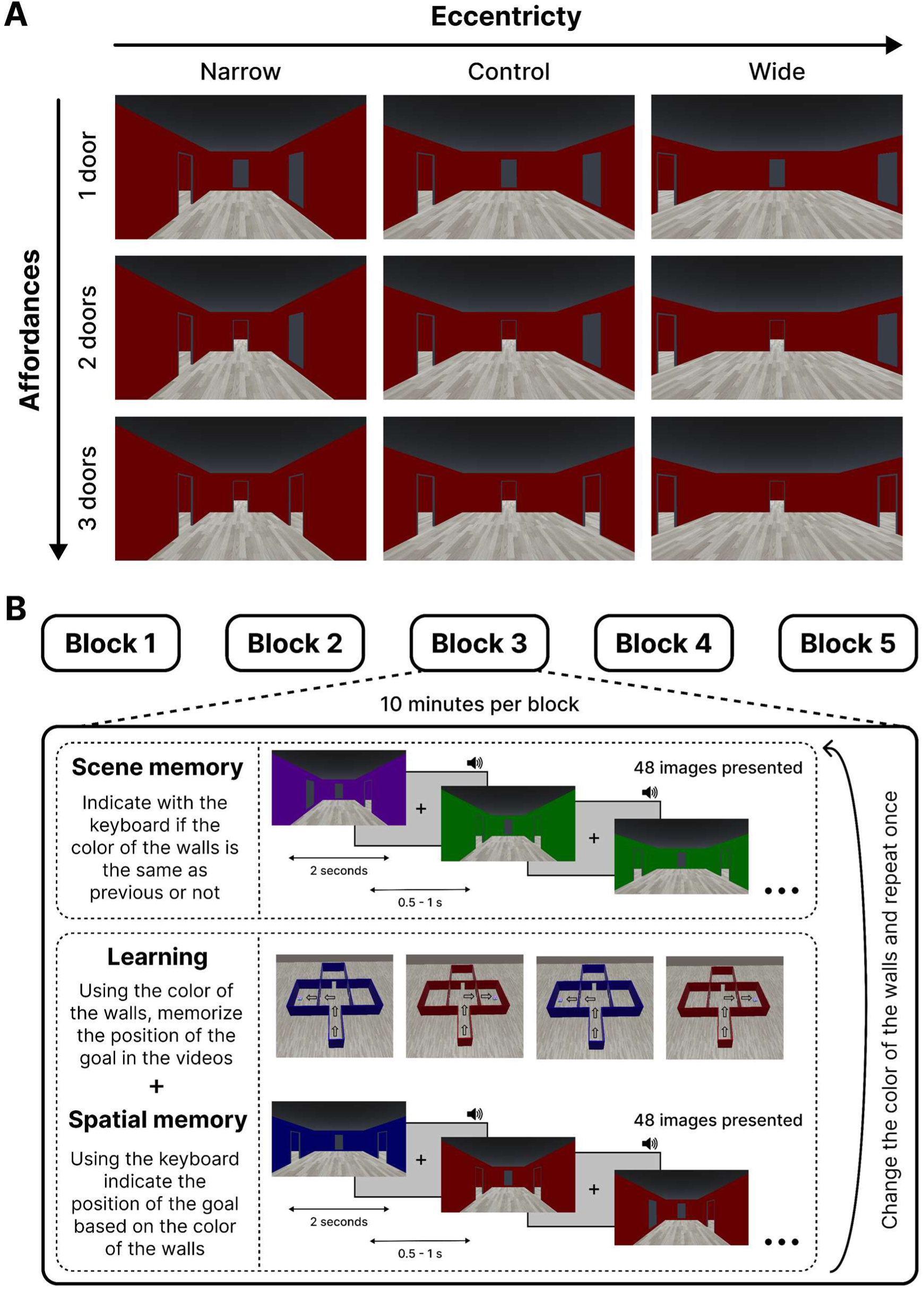
Stimuli and experimental protocol. **A**. Example of the stimulus set for one of the five wall colors used in the experiment, illustrating the different combinations of navigational affordance number and eccentricity. **B**. Overview of the experimental protocol, which consisted of five blocks lasting approximately 10 minutes each. Each block included two repetitions of an alternating sequence of the scene memory and spatial memory tasks, using distinct wall colors. In each block, 48 images were presented for 2 seconds each, with task-relevant wall color and task-irrelevant affordance configuration systematically varied across trials. The order of the scene and spatial memory task was counterbalanced across blocks and participants.

During the experimental session, participants were seated in a dimly lit, sound-attenuated room to minimize external visual and auditory distractions. Visual stimuli were presented on a 26.5-inch ROG Swift OLED PG27AQDM monitor, positioned at a standardized viewing distance of 60 cm. This OLED display offers a resolution of 2560 × 1440 pixels, a refresh rate of 240 Hz, and an ultra-low response time of 0.03 ms, providing high contrast and accurate color reproduction, critical for the precise rendering and timing of visual scenes in cognitive science experiments (Dimigen & Stein, 2024). Stimulus presentation and timing control were handled using PsychoPy (version 2022.2.2; Peirce et al., 2019) running on a Dell Precision 7560 workstation equipped with an Intel Xeon W-11955 processor. Participants responded using a Razer Huntsman V3 Pro keyboard featuring three individually illuminated buttons, allowing for reliable and temporally accurate recording of response times with millisecond resolution.

The experimental protocol consisted of five blocks, each lasting approximately 10 minutes, for a total session duration of around one hour. The blocks were presented in a pseudorandomized order following a Latin square design and each comprised two phases: *a scene memory* phase and a *spatial memory* phase. The order of these phases varied across blocks to ensure counterbalancing across participants. In the scene memory phase, participants performed a 1-back recognition task. They were required to indicate whether the current visual scene was similar to the one immediately preceding it based solely on the color of the walls, using two designated keys on the keyboard to respond “*Yes*” or “*No*”. Images varied between two rooms with different wall colors, each presented in a pseudorandomized order to ensure that the same color was not presented three times in a row. Through trials, we systematically varied the number of navigational affordances available (*i.e.,* doors or grey rectangles) and their eccentricities (narrow, control, and wide conditions). In the spatial memory phase, participants first viewed four short video clips depicting navigation through two rooms with different wall colors, each leading to a goal located in an adjacent room (*i.e.,* left, front, and right). Participants were instructed to memorize the goal locations, with wall color serving as the only reliable contextual cue. After passively watching these videos, they were presented with static scenes and they were asked to retrieve and indicate the spatial location of the goal based on the room’s wall color (*e.g.,* “In the blue room, the goal is behind the left door; in the red room, it is on the right”). Responses were made using the same keys as in the scene memory phase, and participants were similarly encouraged to respond as quickly and accurately as possible. In both phases, each image was presented for a fixed duration of 2 seconds and followed by a fixation cross displayed for a jittered interval between 500 and 1000 milliseconds. A brief auditory feedback signal was provided after each response to inform participants of their accuracy. Within each block, participants were shown 48 unique images per phase, each repeated twice, resulting in 96 trials per phase per block. Across the five blocks, this yielded a total of 480 image presentations per phase. Stimuli were evenly distributed across conditions manipulating the number and eccentricity of navigational affordances, ensuring balanced representation of all experimental conditions. We also ensured that each image always contained at least one open door, corresponding to the door leading to the goal. Critically, in both scene memory and spatial memory phases, only the color of the walls was task-relevant, while the eccentricity and the number of navigational affordances were systematically manipulated but remained irrelevant to task demands.

### Statistical analysis

All the statistical analyses were performed using R Statistical Software (version 4.5.1, R Foundation for Statistical Computing, Vienna, Austria) with R studio (version 2025.05.1). To ensure the robustness and interpretability of the statistical models, while also minimizing computational complexity and avoiding convergence issues, we first averaged reaction times across the different sublevels of experimental conditions: *Age* (young *vs* older), *Phase* (scene memory *vs* spatial memory), *Affordances* (1 door *vs* 2 doors *vs* 3 doors) and *Eccentricity* (narrow *vs* control *vs* wide). This preprocessing step allowed us to reduce trial-level noise and focus on condition-level effects, which were the primary focus of our hypotheses. Given the high level of accuracy observed among participants (young adults: M = 96.53%, SD = 1.91; older adults: M = 92.67%, SD = 4.65), we included only trials with correct responses in subsequent analyses, meaning that participants accurately recalled the position of the goal during the spatial memory phase. To determine the optimal structure of fixed and random effects, we specified and compared a total of 50 candidate models, systematically varying the inclusion of predictors and random structures (see **Supplementary Table 1**). Model comparison was based on the Akaike Information Criterion (AIC), which balances model fit and complexity. The model with the lowest AIC was selected as the best fitting model for subsequent analyses. We confirmed that the selected model was not singular and that model residuals met the assumption of normality based on visual inspection of quantile-quantile (Q-Q) plots. Fixed effects were tested using Type III analysis of variance with Satterthwaite’s approximation for degrees of freedom, as implemented in the *anova()* function from the *lmerTest* package. Effect sizes were estimated using partial eta-squared (η_p_^2^). Estimated Marginal Means (EMMs) were computed using the *emmeans* package, and post-hoc comparisons were performed using Tukey’s Honest Significant Difference (HSD) correction to account for multiple comparisons, with Cohen’s d as measure of effect size.

After comparison with other models (see **Supplementary Table 1**), the following model was retained:

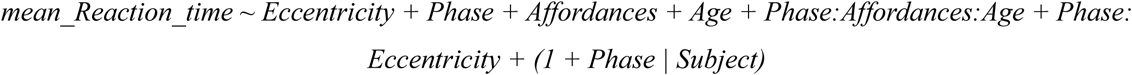

Here we treated the number of affordances as a categorical variable to allow the extraction of the EMMs, before subsequently modeling it as a continuous variable to estimate the slope and enable comparisons between groups and conditions. For the sake of clarity in the results, we initially use the exact terminology of Estimated Marginal Mean (EMM) and Standard Error (SE) with the SE reflecting the precision of the model-based estimate across participants. Later in the text, we simplify the notation by replacing EMM with *M* and reporting SE after “±” to improve readability.

On a related note, we also analyzed accuracy. Given our methodological choices, accuracy was binary at the trial level (correct *vs.* incorrect) and showed a pronounced ceiling effect, which violated the assumption of normality of the residuals made in linear mixed-effects modeling. Accordingly, accuracy was analyzed using a generalized linear mixed-effects model with a binomial distribution and logit link function. This analysis revealed no significant effects except for phase (see **Supplementary 2**). Importantly, the task was deliberately designed to yield high accuracy, maximizing the number of valid responses and minimizing speed–accuracy trade-offs, thereby preserving sensitivity to the reaction-time effects that were the primary focus of the study.

## Results

### Older adults are slower, especially during scene memory phase

First, as expected, older adults were slower at performing the task (Estimated Marginal Means = 0.951, SE = 0.021) compared to young (M = 0.700 ± 0.019; F_(1,50.95)_ = 79.47, *p* < .001, η_p_^2^ = 0.61, 95% CI = [0.44, 0.72]), replicating classical age-related slowdown. Overall, participants were also slower to answer in the scene memory phase (M = 0.880 ± 0.016) compared to the spatial memory phase (M = 0.771 ± 0.014), a difference that was even more pronounced for older adults (M = 1.018 ± 0.020 *vs* M = 0.885 ± 0.020; t_(50)_ = 11.24, *p* < .001, d = 5.71, 95% CI = [4.68, 6.80]), compared to younger (M = 0.657 ± 0.019 *vs* M = 0.743 ± 0.021; t_(50)_ = 7.80, *p* < .001, d = 3.69, 95% CI = [2.72, 4.65]; **Figure 2.A**). This difference likely reflects the 1-back nature of the scene-memory task, which requires continuously updating the representation of the previously seen image with the current one, a control process that deteriorates with age (Oberauer, 2005; Verhaeghen & Basak, 2005).

**Figure 2.**
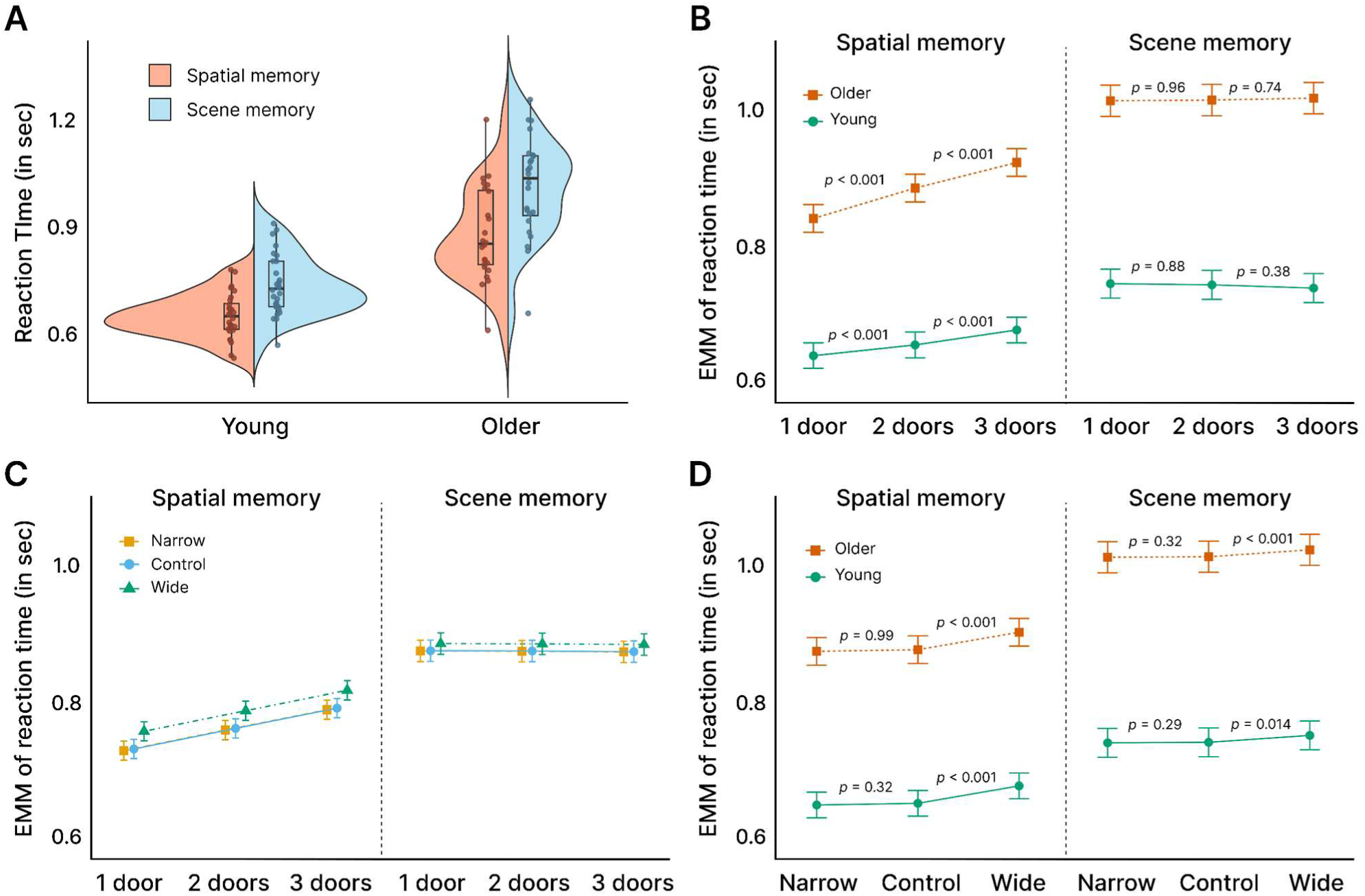
Presentation of the results for the reaction time. **A**. Mean reaction time per subject comparing scene and spatial memory phases across age groups. **B**. Estimated Marginal Means (EMMs) with the Standard Error (SE) for the three-way interaction between the phase, the affordances, and the age. **C**. EMMs with the SE for the three-way interaction between the phase, the condition, and the affordances. No statistical results are presented in this panel, as this interaction was not present in the model selected based on the AIC values. **D**. EMMs with the SE for the three-way interaction between the phase, the condition, and the age. In the selected statistical model, there was no interaction with age, but for graphical purposes we decided to separate the age groups to highlight the similar trend between ages.

To further investigate if this observed difference stems from this updating process, we conducted a follow-up analysis investigating whether visual similarity to the preceding image influenced participants’ reaction times. In each block, two distinct wall colors were presented in a pseudorandom order, ensuring that the same color did not appear more than thrice consecutively. Participants were instructed to detect color changes in the scene memory task. However, in the spatial memory task, although the same color alternation occurred, this feature was not relevant to the task. Analysis of reaction times revealed a significant three-way interaction among phase, age, and similarity of the image to the previous one F_(4,100.93)_ = 84.97, *p* < .001, η_p_^2^ = 0.77, 95% CI = [0.69, 0.82]). Here, for the trials in which the color differed from the previous color, performance decreased for older adults more for the scene memory (M = 1.043 ± 0.023 *vs* M = 0.986 ± 0.023; t_(1758)_ = 14.10, *p* < .001, d = 1.36, 95% CI = [1.16, 1.56]) than for the spatial memory (M = 0.895 ± 0.020 *vs* M = 0.864 ± 0.020; t_(1758)_ = 7.68, *p* < 0.001, d = 0.74, 95% CI = [0.54, 0.93]). For young adults the effect was less pronounced for the scene memory (M = 0.757 ± 0.021 *vs* M = 0.725 ± 0.021; t_(1758)_ = 8.63, *p* < .001, d = 0.77, 95% CI = [0.59, 0.95]) and was not significant for the spatial memory (t_(1758)_ = 1.61, *p* = .38). These results suggest that older adults are particularly vulnerable to disruptions in task-relevant perceptual continuity (*i.e.,* wall color). That is especially relevant here, when participants have to update the information in working memory in response to a new wall color, while simultaneously comparing it to the preceding color to successfully complete the current trial. In other words, for older adults, the *blue–red–blue* sequence was more difficult than the *blue–blue–red* sequence, but only during the scene memory phase, which required remembering the previous color.

### Reaction time increased linearly with the number of navigational affordances during spatial memory task, an effect amplified in older adults

Based on previous evidence for an increased reliance on prior information in older adults, we examined how aging modulates the interaction between bottom-up (*i.e.,* number of affordances) and top-down (*i.e.,* prior spatial knowledge) processing. We reported a very large effect for the interaction between the number of affordances, the phase and the age group (F_(7,146.50)_ = 94.74, *p* < .001, η_p_^2^ = 0.82, 95% CI = [0.77, 0.85]). The overall pattern was similar across age groups, with an increase in reaction time as the number of affordances increased, but only in the spatial memory phase. That is, in the spatial memory phase, reaction time significantly increased with the number of affordances (one: M = 0.741 ± 0.014; two: M = 0.770 ± 0.014; three: *M* = 0.801 ± 0.014) with all pairwise comparisons reaching significance: performance decreased from one to two affordances (*p* < .001, d = 0.43), from one to three (*p* < .001, d = 0.86), and from two to three (*p* < .001, d = 0.42). In contrast, in the scene memory phase, reaction time remained constant across all affordance levels (one: M = 0.881 ± 0.016; two: M = 0.881 ± 0.016; three: M = 0.884 ± 0.016), with no significant differences between conditions (all *p* > .87).

With age, the increase in reaction time associated with a higher number of affordances during the spatial memory phase was even more pronounced. Indeed, in the spatial memory phase, older adults showed a stronger sensitivity to the number of affordances, with significantly larger differences (*e.g.,* one *vs.* two: *p* < .001, *d* = 1.93; one *vs.* three: *p* < .001, *d* = 3.56; two *vs.* three: *p* < .001, *d* = 1.63) compared to younger adults (one *vs.* two: *p* < .001, *d* = 0.69; one *vs.* three: *p* < .001, *d* = 1.64; two *vs.* three: *p* < .001, *d* = 0.95; **Figure 2.B**). Modeling affordances as a continuous variable confirmed a significant positive linear trend in the spatial memory phase for both age groups, with the effect being markedly stronger in older adults. Specifically, in the spatial memory phase, reaction time increased by approximately 40ms per additional affordance (*β* = 0.041, 95% CI [0.036, 0.045]), which was more than twice the change observed for younger adults (*β* = 0.020, 95% CI [0.015, 0.024]). Importantly, the model selected based on AIC did not include an interaction with eccentricity, indicating that adding this term did not improve model fit sufficiently to justify its increased complexity. This suggests that the eccentricity of the open doors did not meaningfully influence the level of interaction between the number of affordances and the phase, a pattern that was confirmed when we extracted and plotted the estimated marginal means (**Figure 2.C**). To further confirm this absence of effect, we forced the selection of a linear mixed-effects model including the full interaction. As expected, we observed no interaction between affordances, eccentricity, and phase (F_(4,884)_ = 1.42, *p* = .23), nor a higher-order interaction including age (affordances × eccentricity × phase × age: F_(4,884)_ = 0.29, *p* = .89). Together, these results confirm that a distractor-based explanation is unlikely to account for the interaction observed between the number of affordances and the phase.

### Scene openness makes participants slower, particularly during the spatial memory phase

Finally, the selected statistical model showed an interaction between the eccentricity of the affordances and the phase, which we investigated further. We firstly reported a main effect of the eccentricity (F_(2,820)_ = 69.03, *p* < .001, η_p_^2^ = 0.14, 95% CI = [0.10, 0.19]), with no difference between the narrow (M = 0.817 ± 0.014) and the control conditions (M = 0.820 ± 0.014; *p* = .653). However, both were different from the wide condition (M = 0.838 ± 0.014; both *p* < .001), with d = 0.85 (95% CI = [0.68, 1.01]) and d = 0.77 (95% CI = [0.61, 0.94]) compared with the narrow and control conditions, respectively. There was a significant interaction between condition and phase (F_(2, 820)_ = 13.18, *p* < .001, η_p_^2^ = 0.03, 95% CI = [0.01, 0.06]), and while the pattern stayed the same, the magnitude of the differences between the conditions was greater in the spatial memory phase. In the spatial memory phase, the wide condition differed from the control (d = 1.11, 95% CI = [0.88, 1.35]) and the narrow conditions (d = 1.22, 95% CI = [0.99, 1.46]). In contrast, in the scene memory phase these differences were much smaller: wide *vs.* control (d = 0.44, 95% CI = [0.21, .67]) and wide *vs.* narrow (d = 0.27, 95% CI = [0.05, 0.70]; **Figure 2.D**). These results suggest that participants required more time to respond as the eccentricity of the affordances increased, and that this slowdown was even more pronounced for the spatial memory phase. However, as reported previously, there was no interaction between eccentricity and the number of affordances, ruling out a distractor-based explanation, which would have predicted such an interaction.

Instead, one possibility is that the observed main effect of eccentricity reflects slower responses when participants interact with a more open scene, a global scene property that is known to influence early neural processing and could predict behavioral response speed (Greene & Oliva, 2009; Hansen et al., 2018; Orima & Motoyoshi, 2023). To dissociate the contribution of door eccentricity from that of overall room openness, we focused on trials with a single door. In these trials, the room geometry was fixed such that the door was in the central position. We then compared trials featuring a single central door (*i.e.,* constant affordance eccentricity while room size increased) with trials in which the single door was placed on a lateral wall (*i.e.,* affordance eccentricity increasing as room size increased). This analysis reported no main effect of the position of the door (F_(1,514.99)_ = 1.70, *p* = .19) and no interaction between the position of the door and the condition room openness (F_(1,514.99)_ = 1.93, *p* = .15), suggesting that the eccentricity of the affordance was not the factor influencing the increased reaction time, but that the openness of the room was the main driver of this increase.

## Discussion

The present behavioral study investigated how aging impacts the interplay between the bottom-up processing of scene features, and the top-down modulation elicited by the acquisition of spatial knowledge about the scene. We showed that reaction times increased linearly with the number of navigational affordances (*i.e.,* open doors) only after participants had learned the spatial location of the goal in an adjacent room, even though the visual stimuli presented were strictly identical before and after learning. Modulating the eccentricity of the open doors did not affect this interaction, reinforcing our results by allowing us to rule out distraction-based explanations, as attentional distraction typically varies with stimulus position in the visual field. Crucially, the magnitude of this interaction was twice as large in older participants compared to young adults. Together, these findings confirm that prior spatial knowledge interact with the processing of visual scene features, and that aging selectively amplifies this interaction, leading to slower interactions with visual scenes features.

### Prior spatial knowledge induces a linear affordance-related performance cost

Firstly, we confirmed that prior spatial information systematically interferes with the extraction of navigational affordances, resulting in a linear decline in performance (Naveilhan et al., 2024). In the current results, this decrease in performance was measured by reaction times, as not decrease in accuracy was observed, mainly due to a ceiling effect (see **Supplementary 2**). This ceiling effect can be explained by the ease of both tasks, amplified by the increase in image presentation time from 1000 ms to 2000 ms compared to the Naveilhan et al. (2024) protocol to account for older adults’ slower responses, and to avoid a potential speed–accuracy trade-off that is modulated with aging. Older adults typically adopt a more conservative response criterion (prioritizing accuracy over speed) and show reduced flexibility in shifting toward emphasizing speed in perceptual decision tasks (Forstmann et al., 2011; Ramanoël et al., 2015; Starns & Ratcliff, 2010). Increasing image presentation time may be an interesting factor to investigate this interaction, because Ménétrier et al. (2018) proposed that longer encoding time may allow higher-level knowledge to modulate or override bottom-up sampling predictions. In our results, the number of lower-level navigational affordances only affects performance once such knowledge is available, pointing to a top-down modulation process rather than simple feedforward distraction. The absence of affordance-related modulation during the scene-memory phase is consistent with broader evidence that core scene properties are extracted rapidly and that early scene-selective responses are strongly constrained by low– and mid-level visual structure before being further shaped by higher-level, goal– and context-dependent interpretations (Groen et al., 2017; Malcolm et al., 2016).

However, it remains possible that during the spatial memory task, the increased reaction times were due to the other affordances acting as distractors. To disambiguate this possibility, we varied the degree of eccentricity of the open doors, since distractor interference is known to vary with eccentricity. Prior studies have shown that peripheral distractors result in greater manual and saccadic reaction time costs than central ones, reflecting weaker inhibitory control at greater eccentricities (Chen, 2008; Chen & Treisman, 2008; Honda, 2005). On the other hand, central distractors may be more disruptive due to privileged foveal access, a pattern further refined by recent EEG evidence showing that perceptual load reduces interference from peripheral but not central distractors (Beck & Lavie, 2005; Chen et al., 2023). While our aim is not to distinguish between these competing explanations that may depend on the context and the nature of the distractors, we emphasize that if the open doors had been consciously attended to (*e.g.,* via visual saccades) or processed solely as distractors, their eccentricity would likely have influenced our results. In the present results, the eccentricity of the open doors did not influence the reported interaction between affordance extraction and prior spatial information (**Figure 2.C**). Although this absence of modulation strongly argues against a distractor-based explanation, it cannot be entirely ruled out. During the spatial memory phase, when participants are required to retrieve the goal location, they may still consider alternative action possibilities in the scene (even when the only task-relevant feature is the wall color) by actively visually sampling these affordances within an orienting task context. While this possibility remains unlikely given evidence that representations of navigational affordances are robust across task demands (Bartnik et al., 2025), future work incorporating eye-tracking measures could directly test this possibility by determining whether patterns of visual exploration differ across task phases. Taken together, the replication of previous findings in a new, independent dataset strengthens our earlier evidence that prior spatial information interacts with the extraction of navigational affordances (Naveilhan et al., 2024, 2025). This result is further supported by the observation that the modulation is unaffected by affordance eccentricity and does not occur before participants acquire the relevant spatial priors, despite identical stimulus presentations in the scene memory and spatial memory phases.

### Aging amplifies the interaction between prior knowledge and the extraction of scene features

The present study also aimed to examine whether the modulation by prior spatial information extends to older adults. Growing literature suggests that aging is associated with a shift toward greater reliance on top-down modulation from previous knowledge when bottom-up processing becomes less reliable, slower or more noisy (Borges et al., 2020; Lai et al., 2020; Musel et al., 2012; Neider & Kramer, 2011; Power & Conlon, 2017; Ramzaoui et al., 2021). These elements suggest two opposing explanations regarding the role of top-down modulation in aging. One interpretation is that older adults increasingly depend on top-down spatial expectations, which may lead them to disregard the actual number of affordances presented. An alternative interpretation is that the influence of prior spatial information may be more strongly weighted in older individuals, thereby amplifying the interaction between spatial context and affordance extraction. Our results support the latter, with older adults displaying an increased interaction effect twice as large as the one reported for young adults. Thus, older adults appear to rely more heavily on acquired spatial priors when extracting navigational affordances, a pattern consistent with the view that acquired knowledge can compensate for age-related memory limitations, while also increasing susceptibility to knowledge-driven costs when it biases or constrains ongoing processing (Umanath & Marsh, 2014). This age-related heightened weighting of spatial priors can account for the steeper decline in performance observed in older participants. Evidence from previous research supports this interpretation: both facilitative and detrimental effects of prior cues are more pronounced in older adults, indicating a greater dependence on such information to guide visual search strategies (Neider & Kramer, 2011). In the same vein, Lai et al. (2020) found that healthy older adults showed behavioral sensitivity to scene congruity and reported a delayed and enlarged N1 component consistent with degraded early sensory input, with older adults also showing larger P2 for blurred targets. This last result was interpreted as an early deployment of top-down resources to resolve perceptual ambiguity. Thus, in future work it would be interesting to investigate whether the effect of age reported in the present study is also reflected at the neural level, for instance by examining the modulation of the P2 component over occipito-parietal electrodes (Harel et al., 2022; Naveilhan et al., 2024).

Finally, our findings may also be interpreted in light of the *environmental support* hypothesis of cognitive aging (for a review, see Lindenberger & Mayr, 2014). Converging evidence indicates that older adults could also exhibit an over-reliance on bottom-up information from the environment to compensate for age-related declines in core cognitive functions such as attention, memory, and executive control. In line with this view, increased reliance on low-level environmental information has been associated with reduced engagement of higher-order task processes and diminished top-down modulation in older adults during goal-directed behavior (Mayr, 2001). In this context, the greater increase in reaction times as a function of the number of affordances observed in older adults could be interpreted as an age-related over-reliance on environmental information conveyed by visual scene features. However, under this hypothesis, older adults should have shown a comparable modulation of reaction time by the number of affordances during the scene-memory phase, which was not observed here. To definitively rule out this hypothesis, future studies could introduce incongruent trials (*e.g*., the only open door located on the left while the goal is on the right). Under such conditions, the environmental support hypothesis would predict higher error rates in older adults, reflecting a tendency to select the salient but misleading affordance information, or longer reaction time due to the need to inhibit the prepotent affordance-driven response in favor of the task-relevant one.

To conclude, our findings demonstrate that previously acquired spatial knowledge about the goal direction interacts with the extraction of navigational affordances from visual scenes. This top-down modulation emerges only after learning and is robust to eccentricity variations in the visual positioning of affordances. Crucially, this interaction was markedly amplified in older adults, whose performance showed a steeper decline with an increasing number of affordances. These results align with the broader literature suggesting that aging enhances reliance on top-down modulations to compensate for perceptual decline. By showing that prior knowledge not only supports but might also interfere with visual scene processing in aging, this study underscores the dynamic interplay between memory and perception across the lifespan. Thus, while prior spatial information can guide perception, it can also slow older adults’ interactions with complex scenes, an insight that can inform the design of environments and interfaces to reduce these costs while preserving the benefits of prior knowledge.

## Data availability

The complete dataset, including all generated stimuli and analysis code, is publicly available on the OSF repository: https://osf.io/vsg2a/?view_only=56d56acf35b0493b84b3637f7bc2f8bd

## Author Contributions

**Clément Naveilhan**: Conceptualization; Formal analysis; Investigation; Methodology; Writing-Original draft; Writing-Review & editing. **Raphaël Zory:** Funding acquisition; Writing-Review & editing. **Stephen Ramanoël**: Conceptualization; Funding Acquisition; Methodology; Project administration; Supervision, Writing-Review & editing.

## Acknowledgements

This research was made possible thanks to the contribution of our volunteer participants, whose time and engagement are sincerely appreciated by the authors.

This research was supported by the National Research Agency (ANR-25-CE28-7352). This work was also supported by the French government through the France 2030 investment plan managed by ANR, as part of the Initiative of Excellence Université Côte d’Azur under reference number ANR-15-IDEX-01 and, in particular, by the interdisciplinary Institute for Modeling in Neuroscience and Cognition (NeuroMod) of Université Côte d’Azur.

## Supplementary

**Supplementary Table 1.** Comparison of the different statistical models and their associated AIC values. The model highlighted in red was selected based on the lowest AIC, with the additional criterion of normally distributed residuals.

| Rank | Model | AIC |
| --- | --- | --- |
| 1 | mean_Reaction_time ~ Condition + Phase + Affordances + Age + Phase:Affordances:Age + Phase:Condition + (1 + Phase \| Subject) | -3743.41 |
| 2 | mean_Reaction_time ~ Condition + Phase + Affordances + Age + Phase:Affordances:Age + (1 + Phase \| Subject) | -3740.39 |
| 3 | mean_Reaction_time ~ Condition + Phase + Affordances + Age + Phase:Affordances:Age + Phase:Condition + Condition:Affordances + (1 + Phase \| Subject) | -3732.82 |
| 4 | mean_Reaction_time ~ Condition + Phase + Affordances + Age + Phase:Affordances:Age + Phase:Condition + Condition:Age + (1 + Phase \| Subject) | -3726.15 |
| 5 | mean_Reaction_time ~ Condition + Phase + Affordances + Age + Phase:Affordances + Age:Condition + (1 + Phase \| Subject) | -3693.42 |
| 6 | mean_Reaction_time ~ Condition * Phase * Affordances + Age + (1 + Phase \| Subject) | -3677.36 |
| 7 | mean_Reaction_time ~ Condition * Phase * Affordances * Age + (1 + Phase \| Subject) | -3613.24 |
| 8 | mean_Reaction_time ~ Condition + Phase + Affordances + Age + (1 + Phase + Age \| Subject) | -3535.74 |
| 9 | mean_Reaction_time ~ Condition + Phase + Affordances + Age + Condition:Phase + (1 + Phase + Age \| Subject) | -3532.02 |
| 10 | mean_Reaction_time ~ Condition + Phase + Affordances + Age + (1 + Phase + Condition + Affordances + Age \| Subject) | -3527.57 |
| 11 | mean_Reaction_time ~ Condition + Phase + Affordances + Age + (1 + Phase \| Subject) | -3526.49 |
| 12 | mean_Reaction_time ~ Condition + Phase + Affordances + Age + (1 \| Subject/Phase) | -3525.87 |
| 13 | mean_Reaction_time ~ Condition + Phase + Affordances + Age + Condition:Phase + (1 + Phase \| Subject) | -3522.77 |
| 14 | mean_Reaction_time ~ Condition + Phase + Affordances + Age + Condition:Phase + (1 \| Subject/Phase) | -3522.16 |
| 15 | mean_Reaction_time ~ Condition + Phase + Affordances + Age + Phase:Condition + (1 + Phase \| \| Subject) | -3520.77 |
| 16 | mean_Reaction_time ~ Condition + Phase + Affordances + Age + Condition:Phase + Condition:Affordances + (1 + Phase + Age \| Subject) | -3512.74 |
| 17 | mean_Reaction_time ~ Condition + Phase + Affordances + Age + Condition:Phase + Condition:Affordances + (1 + Phase + Condition + Affordances + Age \| Subject) | -3508.57 |
| 18 | mean_Reaction_time ~ Condition + Phase + Affordances + Age + Affordances:Condition + (1 + Phase \| Subject) | -3506.8 |
| 19 | mean_Reaction_time ~ Condition + Phase + Affordances + Age + Condition:Phase + Condition:Affordances + (1 + Phase \| Subject) | -3503.49 |
| 20 | mean_Reaction_time ~ Condition + Phase + Affordances + Age + Condition:Phase + Condition:Affordances + (1 \| Subject/Phase) | -3502.88 |
| 21 | mean_Reaction_time ~ Condition + Phase + Affordances + Age + Condition:Phase:Age + (1 + Phase + Age \| Subject) | -3495.15 |
| 22 | mean_Reaction_time ~ Condition + Phase + Affordances + Age + (1 + Age \| Subject) | -2978.26 |
| 23 | mean_Reaction_time ~ Condition + Phase + Affordances + Age + (1 \| Subject) | -2973.3 |
| 24 | mean_Reaction_time ~ Condition + Phase + Affordances + Age + (1 \| Subject) + (1 \| Phase) | -2971.3 |
| 25 | mean_Reaction_time ~ Condition + Phase + Affordances + Age + Condition:Phase + (1 + Age \| Subject) | -2965.8 |
| 26 | mean_Reaction_time ~ Condition + Phase + Affordances + Age + Condition:Phase + (1 \| Subject) | -2960.85 |
| 27 | mean_Reaction_time ~ Condition + Phase + Affordances + Age + Condition:Phase + (1 \| Subject) + (1 \| Phase) | -2958.85 |
| 28 | mean_Reaction_time ~ Condition + Phase + Affordances + Age + Condition:Phase + Condition:Affordances + (1 + Age \| Subject) | -2936.1 |
| 29 | mean_Reaction_time ~ Condition + Phase + Affordances + Age + Condition:Phase + Condition:Affordances + (1 \| Subject) | -2931.14 |
| NA | mean_Reaction_time ~ Condition + Phase + Affordances + Age + (1 + Condition \| Subject) | Singular |
| NA | mean_Reaction_time ~ Condition + Phase + Affordances + Age + (1 + Affordances \| Subject) | Singular |
| NA | mean_Reaction_time ~ Condition + Phase + Affordances + Age + (1 + Phase + Condition \| Subject) | Singular |
| NA | mean_Reaction_time ~ Condition + Phase + Affordances + Age + (1 + Phase + Affordances \| Subject) | Singular |
| NA | mean_Reaction_time ~ Condition + Phase + Affordances + Age + (1 + Phase + Condition + Affordances \| Subject) | Singular |
| NA | mean_Reaction_time ~ Condition + Phase + Affordances + Age + Condition:Phase + (1 + Condition \| Subject) | Singular |
| NA | mean_Reaction_time ~ Condition + Phase + Affordances + Age + Condition:Phase + (1 + Affordances \| Subject) | Singular |
| NA | mean_Reaction_time ~ Condition + Phase + Affordances + Age + Condition:Phase + (1 + Phase + Condition \| Subject) | Singular |
| NA | mean_Reaction_time ~ Condition + Phase + Affordances + Age + Condition:Phase + (1 + Phase + Affordances \| Subject) | Singular |
| NA | mean_Reaction_time ~ Condition + Phase + Affordances + Age + Condition:Phase + (1 + Phase + Condition + Affordances \| Subject) | Singular |
| NA | mean_Reaction_time ~ Condition + Phase + Affordances + Age + Condition:Phase + (1 + Phase + Condition + Affordances + Age \| Subject) | Singular |
| NA | mean_Reaction_time ~ Condition + Phase + Affordances + Age + Condition:Phase + Condition:Affordances + (1 + Condition \| Subject) | Singular |
| NA | mean_Reaction_time ~ Condition + Phase + Affordances + Age + Condition:Phase + Condition:Affordances + (1 + Affordances \| Subject) | Singular |
| NA | mean_Reaction_time ~ Condition + Phase + Affordances + Age + Condition:Phase + Condition:Affordances + (1 + Phase + Condition \| Subject) | Singular |
| NA | mean_Reaction_time ~ Condition + Phase + Affordances + Age + Condition:Phase + Condition:Affordances + (1 + Phase + Affordances \| Subject) | Singular |
| NA | mean_Reaction_time ~ Condition + Phase + Affordances + Age + Condition:Phase + Condition:Affordances + (1 + Phase + Condition + Affordances \| Subject) | Singular |
| NA | mean_Reaction_time ~ Condition + Phase + Affordances + Age + Phase:Affordances:Age + Phase:Condition + (1 + Phase + Affordances \| Subject) | Singular |
| NA | mean_Reaction_time ~ Condition + Phase + Affordances + Age + Condition:Affordances + Phase:Age + (1 + Condition \| Subject) | Singular |
| NA | mean_Reaction_time ~ Condition + Phase + Affordances + Age + Affordances:Condition + Phase:Age + (1 + Affordances \| Subject) | Singular |
| NA | mean_Reaction_time ~ Condition * Affordances * Age + Phase + (1 + Affordances + Age \| Subject) | Singular |
| NA | mean_Reaction_time ~ Condition + Phase + Affordances + Age + Affordances:Age + Condition:Phase + (1 + Phase + Affordances \| Subject) | Singular |

## Supplementary 2

In this supplementary analysis we investigated the accuracy, using a generalized linear mixed-effects model due to ceiling effects of this metric (see **Supplementary figure 1**). Results of the GLMM indicated only an effect for the phase (χ^2^_(1)_ = 22.82, *p* < .001) with more errors made during the scene memory phase. There was no effect for the age (χ^2^_(1)_ = 0.20, *p* = .66), the eccentricity (χ^2^_(2)_ = 1.51, *p* = .47), or the affordances (χ²_(1)_ = 1.82, *p* = .40), and no other interactions (all *p* > .078). This lack of effects on accuracy is likely attributable to the pronounced ceiling of performance, reflecting the relative ease of the task and the relatively long response window of 2000 ms. As stated in the Method section, this design choice was intentional, as it maximized the number of correct responses while minimizing speed–accuracy trade-offs, thereby preserving sensitivity to the reaction time effects that constituted the primary focus of the study.

**Supplementary figure 1.**
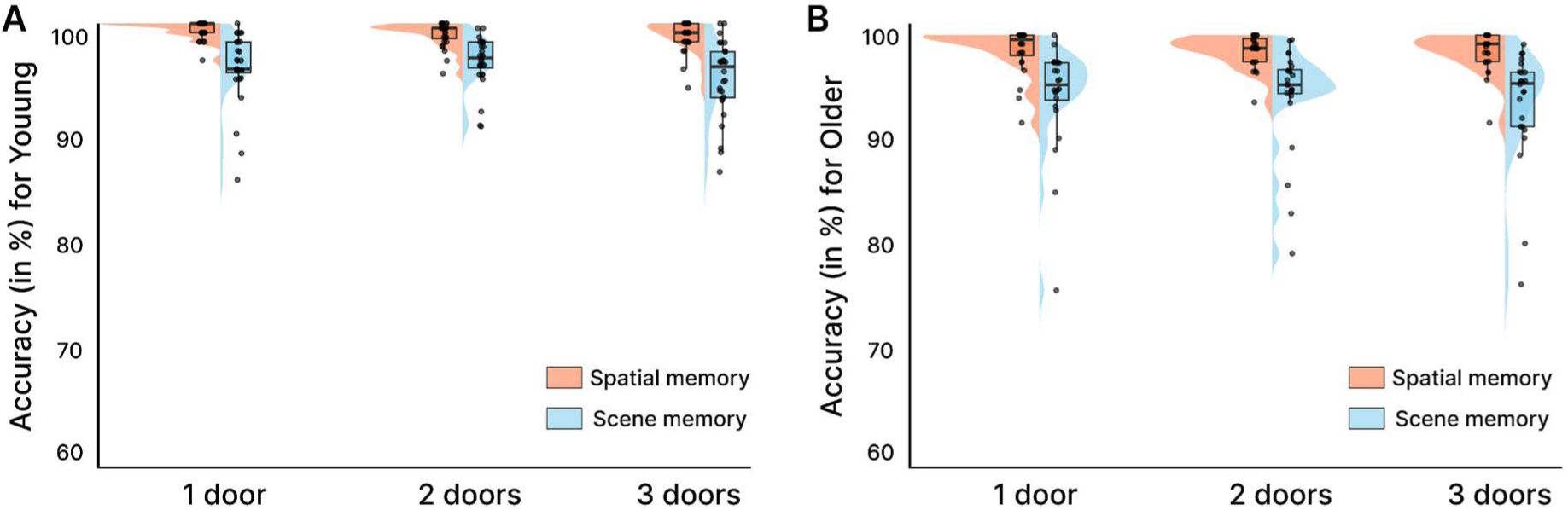
Presentation of the mean values for the accuracy, across phase (scene memory vs spatial memory) and affordances. A. For young adults B. For older adults

